# Ancestral variation and its impact on wild house mouse genomes

**DOI:** 10.1101/2023.11.09.566486

**Authors:** Raman Akinyanju Lawal, Beth L. Dumont

## Abstract

Ancestral alleles are important contributors to adaptation and disease risk in populations. House mice emerged in and/or around the Indian subcontinent, but the genetic composition of this ancestral population and the extent to which ancestral variants contribute to contemporary global mouse diversity are poorly understood. To address these knowledge gaps, we assessed the origins and demographic patterning of global mouse diversity using a set of 169 wild mouse genome sequences from across the world. This dataset includes 37 mouse genomes from the broadly designated ancestral regions, providing crucial resources needed to evaluate the contributions and the impact of ancestral diversity on the genomic scale. We show that house mice emerged in the Indo-Pakistan region around 700 kya, with *M. m. castaneus* at the root of the *M. musculus* species. Migration out of the Indo-Pakistan homeland led to the subsequent emergence of the *M. m. domesticus* and *M. m. musculus* subspecies ∼360 and 260 kya, respectively. A modest fraction of ancestral alleles have persisted long-term across mouse populations through balancing selection, and we demonstrate that such regions are strongly enriched for genes with immune-related functions. Finally, we find widespread allele-sharing across *Mus musculus* lineages and show that this trend is largely due to incomplete lineage sorting, an interpretation contrary to some recent claims of pervasive subspecies introgression. Together, our work underscores the contributions of ancestral variants to contemporary house mouse diversity and adaptation, and refines our understanding of the natural evolutionary history of this important model species.

## INTRODUCTION

House mice (*Mus musculus*) are the premier biomedical model system and a powerful natural system for investigating the genetic and mechanistic basis of adaptation to diverse environments (Boursot, et al. 1993; Phifer-Rixey and Nachman 2015; Lawal, et al. 2021; Beckman, et al. 2022). House mice emerged from a common ancestor in and/or around the Indian subcontinent less than 3 mya (Suzuki, et al. 2004), although the specific ancestral location within this broad geographic region remains uncertain (Boursot, et al. 1993; Boursot, et al. 1996; Din, et al. 1996; Suzuki, et al. 2013). Prior work has suggested that house mice expanded out of this ancestral region approximately ∼150 kya to 500 kya, giving rise to three core subspecies (Boursot, et al. 1996; Geraldes, et al. 2008; Geraldes, et al. 2011; Bonhomme and Searle 2012; Suzuki, et al. 2013; Phifer-Rixey, et al. 2020). *M. m. domesticus* is native to Western Europe and the Iranian Valley.

*M. m. musculus* is present across Eastern Europe and Siberia, and *M. m. castaneus* is distributed across Southeast Asia. Aided by human dispersal routes in recent history, house mice have expanded their footprint outside of these native ranges, colonizing all major continents except for Antarctica, and inhabiting a remarkably broad range of habitats and climates.

The long-term success of house mice in varied environments across the globe is due, at least in part, to their incredible adaptive potential (Lawal, et al. 2021). However, whether house mouse adaptation was largely enabled by selection on new mutations or facilitated by selection on inherited, ancestral alleles remains unknown. A rigorous resolution to this open question relies on a thorough understanding of ancestral mouse genetic diversity and its relationship to genetic variation in contemporary mouse populations. However, there is limited genomic data from mice sampled from the *M. musculus* ancestral range (Harr, et al. 2016; Lawal, et al. 2022) and few population genomic investigations of global wild mouse diversity (Fujiwara, et al. 2022). These resource limitations and standing knowledge gaps also continue to hamper efforts to contextualize the genetic diversity observed in derived (non-ancestral) mouse populations and to accurately infer the demographic and phylogenetic history of wild house mice.

Prior studies of house mice demographic history have relied on sampling small numbers of loci across the genome (Geraldes et al., 2008; Phifer-Rixey et al., 2012) or utilized coalescent strategies dependent on whole genome sequences from a single representative individual for a population or subspecies (Geraldes, et al. 2008; Phifer-Rixey, et al. 2012; Phifer-Rixey, et al. 2020; Fujiwara, et al. 2022). The reliance on a modest number of loci to inform demographic inference is problematic, as few independent lineages coalesce deep in time at any one locus, offering limited power to resolve evolutionary history. Similarly, a restricted number of genomic loci may not faithfully reflect the underlying history of a population, especially if loci are not randomly ascertained. At the same time, demographic inference from a single genome is known to be misleading in the face of population structure or gene flow (Mazet, et al. 2016). These limitations are especially problematic for house mice, which often organize into highly structured demes in nature (Morgan, et al. 2022) and may experience high rates of gene flow between populations (Teeter, et al. 2008; Phifer-Rixey, et al. 2020). Further, single genome inference methods cannot yield accurate estimates of population size in the recent past (Schiffels and Durbin 2014; Terhorst, et al. 2017). This presents a notable shortcoming for demographic inference in house mice, which have undergone human-mediated dispersal across the globe in very recent history. Thus, the most accurate picture of house mouse demographic history stands to be obtained using inference methods that can be applied to multiple whole genomes (Mazet, et al. 2016; Terhorst, et al. 2017).

Recently, we generated whole genome sequences from 14 mice sampled from Pakistan, a location within the broad *M. musculus* ancestral region (Lawal, et al. 2022). Here, we combine this new population genome sequence resource with publicly available genome sequences from wild mouse populations sampled from additional regions within the likely ancestral home range of *Mus musculus* (India, Iran, Afghanistan). We contrast genomic variation in these putative ancestral populations with that found in multiple derived mouse populations sampled from each of the three principal house mouse subspecies (*M. m. domesticus* (DOM), *M. m. castaneus* (CAS), and *M. m. musculus* (MUS)). In total, we analyzed 169 wild mouse genomes including five populations from DOM (America (AMR), France (FRA), Germany (GER), Heligoland (HEL), and Iran (IRA)), three populations from CAS (India (IND), Taiwan (TAI), and Pakistan (PAK)), three populations from MUS (Afghanistan (AFG), Czech Republic (CZR), and Kazakhstan (KAZ)), and the outgroup species, *M. spretus* (SPR). Collectively, our analyses address three fundamental and specific objectives: (1) to re-assess the demographic history of the *M. musculus* lineage and reconstruct the speciation history of the species using the most comprehensive set of wild mouse sequences from the presumed ancestral region to date; (2) to refine the ancestral geographic origins of house mice within the Indian subcontinent and infer the genetic composition of this ancestral population; and (3) to assess the contribution of ancestral genetic diversity to contemporary mouse populations, including an initial exploration of the role of ancestral variation in adaptive evolution. Overall, our work refines current understanding of mouse evolutionary history and spotlights the contribution of ancestral variation to contemporary patterns of genetic diversity in the wild.

## RESULTS

### Subspecies demographic history inferred from wild *M. musculus* genomes

Prior analyses of house mouse subspecies demography have been dependent on limited numbers of genomic loci or single representative genomes (Phifer-Rixey, et al. 2020; Fujiwara, et al. 2022). Here, we adopt an alternative strategy using the scalable approach implemented in SMC++ (Terhorst, et al. 2017) to infer major trends in subspecies demographic history using data from 89 diverse wild mouse genomes representing each of the three principal house mouse subspecies (see Methods). Our analysis yields several key findings.

First, the modern-day CAS subspecies emerged from an ancestral *M. musculus* population around 700 kya with an initial effective population size (*N*_e_) of ∼231,000 (Figure 1a; Table S1). CAS then experienced two subsequent rounds of population expansion and decline. Between 500 kya and 400 kya, the CAS subspecies expanded to *N*_e_ ∼280,000 before experiencing a modest bottleneck to *N*_e_ ∼236,000 at 345 kya. A new wave of expansion began ∼260 kya, lasting until 129 kya (peak *N*_e_ ∼462,000). This expansion was again followed by a population decline to *N*_e_ ∼386,000 around 100 kya. Over the last 36 kya, CAS has maintained a constant *N*_e_ of ∼396,000.

**Figure 1:**
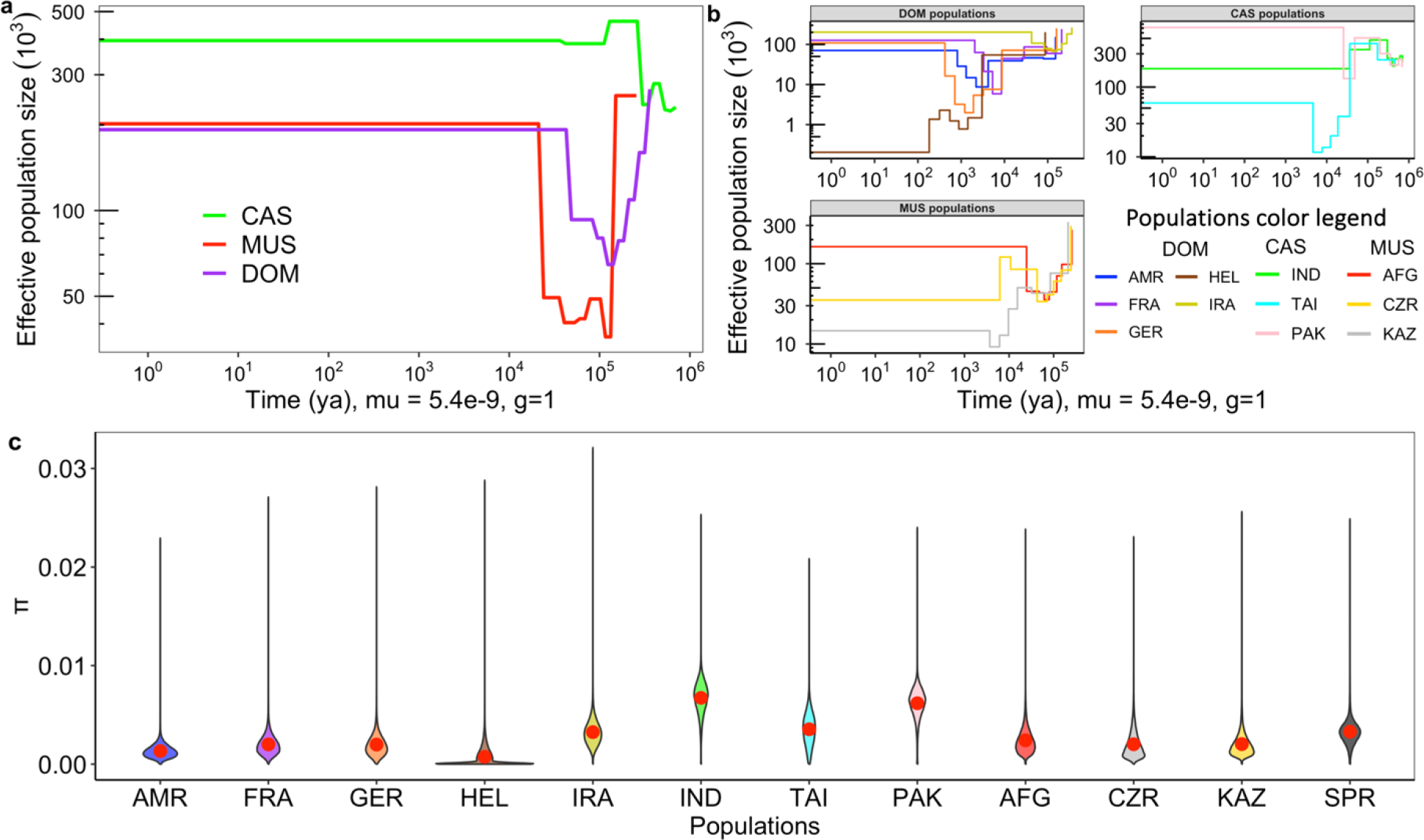
Inferred demographic history for wild house mouse (**a**) subspecies and (**b**) populations using SMC++. Plots depict variation in the effective population size (*N*_e_) as a function of time in years ago (ya). Histories were inferred under the assumptions of 1 generation (g) per year and a per base mutation rate (*μ*) of 5.4x10^-9^. (**c**) Violin plots of the distribution of nucleotide diversity (π) in 100kb windows (50kb slide) in each wild mouse population. The red dot in the middle of each plot corresponds to the genome-wide mean value.

The emergence of the DOM lineage coincided with the first CAS bottleneck ∼ 360 kya (estimated starting *N*_e_ of ∼267,000; Figure 1a). The DOM subspecies then went through a steep population decline, reaching a nadir around 124 kya (*N*_e_ ∼ 65,000). Following a period of recovery, DOM settled at its current *N*_e_ of ∼192,000 at approximately 43 kya.

The MUS lineage emerged ∼256 kya with an initial *N*_e_ of ∼253,000, coincident with the start of the second CAS expansion wave (Figure 1a). Interestingly, the first major bottleneck in MUS (∼130 kya; lowest *N*_e_ ∼36,000) occurred nearly simultaneously with the steep DOM population decline. We speculate that this may coincide with the time of dispersion of both subspecies from their ancestral geographic center. MUS experienced a short expansion between ∼102 kya (*N*_e_ ∼ 49,000) followed by a mild population bottleneck ∼69 kya (*N*_e_ ∼ 42,000) before settling to its current *N*_e_ of ∼202,000 at ∼21 kya (Figure 1a; Table S1).

Our demographic analyses of *M. musculus* subspecies reinforce the presumed ancestral identity of the CAS subspecies. This subspecies was the first *M. musculus* subspecies to diverge from the common ancestral population (Figure S1a) and exhibits the largest historical *N*_e_ — almost twice that estimated for DOM and MUS (Figure 1a). Remarkably, despite differences in methodology, parameter specification (e.g., mutation rate and generation times), and sample data (e.g., sample size, sample source, and number of surveyed sites in the genome), our inferred subspecies split times, are broadly similar to prior published divergence estimates, which span ∼150 - 500 kya; (Boursot, et al. 1996; Geraldes, et al. 2008; Geraldes, et al. 2011; Bonhomme and Searle 2012; Suzuki, et al. 2013; Phifer-Rixey, et al. 2020; Jing Sr, et al. 2023). However, whereas other studies have suggested near simultaneous radiation of the three principal house mouse subspecies [e.g., (Geraldes, et al. 2008; Didion and de Villena 2013; Suzuki, et al. 2013; Phifer-Rixey, et al. 2020)], our analysis discretizes these divergence events in time, suggesting that DOM diverged from the ancestral CAS population approximately ∼100 kya before the MUS subspecies.

**Figure S1:**
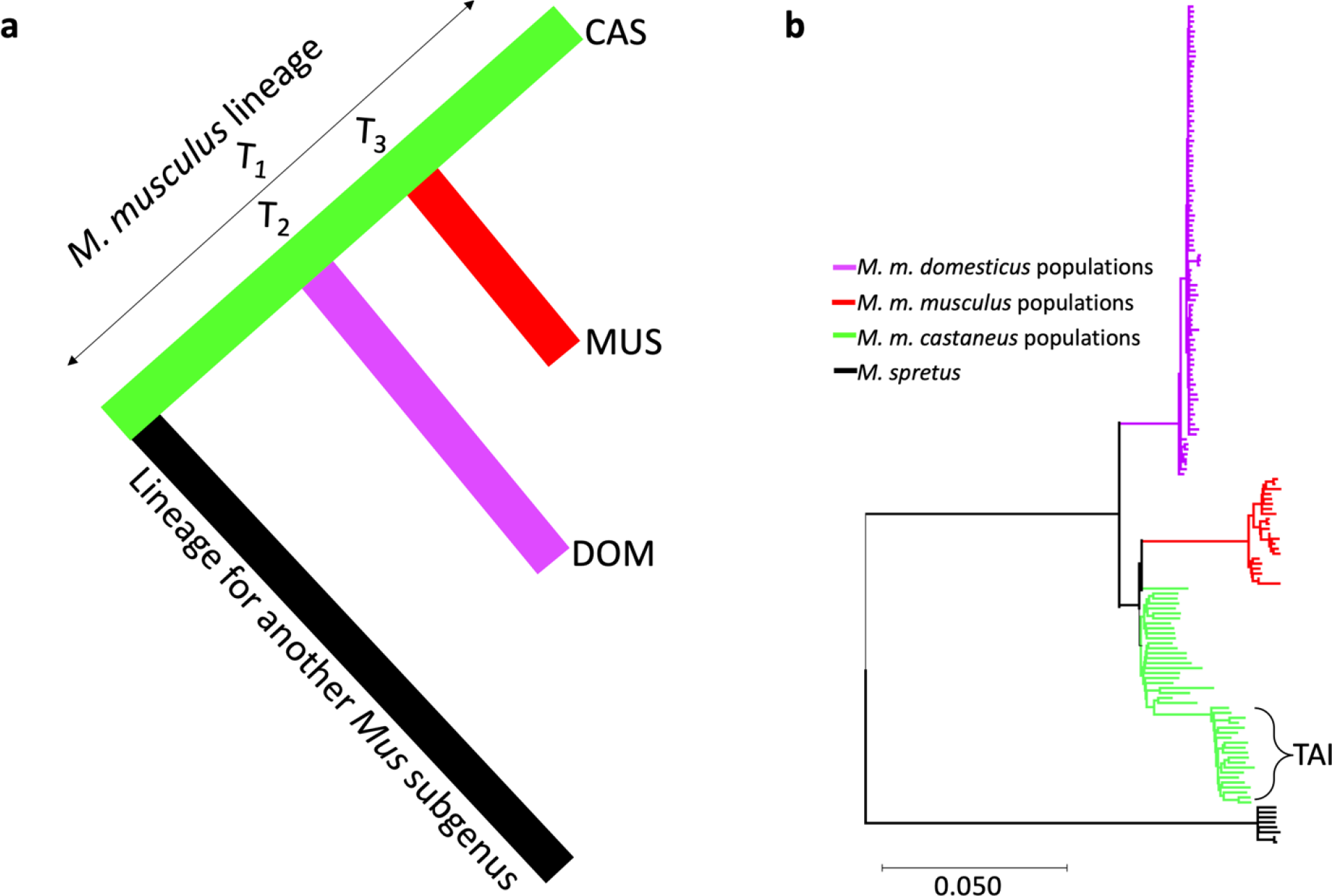
(**a**) Inferred subspecies tree consolidated from the demographic history estimates (see Figure 1a). T_1_ is the time CAS diverged from the common ancestral population of *Mus musculus*, around 700 kya. T_2_ is the DOM divergence time (360 kya) and T_3_ is the MUS divergence time (260 kya). (**b**) Maximum likelihood phylogenetic tree constructed from autosomal SNPs. The Indo-Pakistani populations of CAS are situated at the base of the subspecies tree, consistent with the ancestral identity of these populations. The Taiwanese (TAI) population of CAS is nested within the diversity of the Indo-Pakistan ancestral population.

### Population demographic history inference

We next used SMC++ to infer the history of population size changes across each of the 11 house mouse populations included in our genomic sequence dataset (Figure 1b). While all populations have experienced historical bottlenecks and expansions, population size changes have been most dramatic for *M. m. domesticus* populations from Europe. These observations reinforce conclusions based on limited numbers of genetic markers in European mice, and likely reflect dispersal bottlenecks associated with human-aided transport during the Neolithic and Iron ages, followed by population expansions into new commensal niches in these areas (Bonhomme, et al. 2011; Gabriel, et al. 2011; Bonhomme and Searle 2012). For example, the DOM populations from Germany (GER) and France (FRA) experienced a strong population size contraction ∼8000 ya, roughly coinciding with the start of the Neolithic expansion of human agricultural societies through Europe (Shennan 2018).

The Heligoland population (HEL) experienced an especially dramatic and recent bottleneck (Figure 1b). Mice from this island population were presumably derived through a colonization event from neighboring Germany (GER) (Babiker and Tautz 2015). Indeed, mice from both populations show similar demographic histories until ∼600 ya (Figure 1b; Table S2). However, whereas the GER population spiked in population size at this time, the HEL population plummeted to just a few hundred individuals. While demographic inference is known to be error-prone in the very recent past, the coincidence of the timing of the inferred population bottleneck with the establishment of regular seafaring traffic between Heligoland and Denmark in the 12^th^ century provides orthogonal support for these conclusions (Babiker and Tautz 2015).

### Identifying the geographic origins of *M. musculus*

Contemporary organisms from the ancestral species range can be used as representatives of the ancestral source population (Kirch, et al. 2021). House mice are known to have evolved from an ancestral population in and/or around the Indian subcontinent, but the precise geographic origins of this ancestral population remain subject to dispute (Boursot, et al. 1996; Suzuki, et al. 2013; Hardouin, et al. 2015). Ancestral populations are expected to (1) harbor elevated effective population size, (2) exhibit high genetic diversity, and (3) should largely subsume the genetic variation present in derived populations (Campbell and Tishkoff 2008). We addressed each of these testable predictions by comparing the genomes of the four Indo-Iranian house mouse populations from the putative ancestral regions – IND, PAK, AFG, and IRA – to mice from other global populations.

First, we expected that the ancestral house mouse population should exhibit a large *N*_e_ (Garrigan, et al. 2007; Kapopoulou, et al. 2018). Compared to other surveyed mouse populations, *N*_e_ is the largest of the four populations from Indo-Iranian valley. Of these candidate ancestral populations, PAK has the highest *N*_e_ (∼717,000), followed by IRA (∼202,000), IND (∼184,000), and AFG (163,000) (Figure 1b; Table S2).

Second, contemporary populations residing within the ancestral species home range should harbor greater genetic diversity than mice from non-ancestral, derived populations, as observed in other taxa (Beck, et al. 2008; Campbell and Tishkoff 2008; Pool, et al. 2012). Nucleotide diversity in PAK (0.0062 ± 0.016) and IND (0.0067 ± 0.018) is twice that in IRA (0.0032 ± 0.014), three times that in AFG (0.0024 ± 0.0014), and up to 8 times the level of nucleotide diversity observed in other global mouse populations (Figure 1c). Similarly, PAK and IND harbor twice as many variants compared to IRA and AFG, and up to 14 times that observed in other global populations (Figure 2a). IND and PAK also harbor the greatest numbers of population-private alleles (Figure 2b). Assuming that geographic sampling schema does not bias nucleotide diversity estimates (Blair 1953) and the absence of extreme demographic events, these results appear to exclude IRA and AFG as putative founder populations.

**Figure 2:**
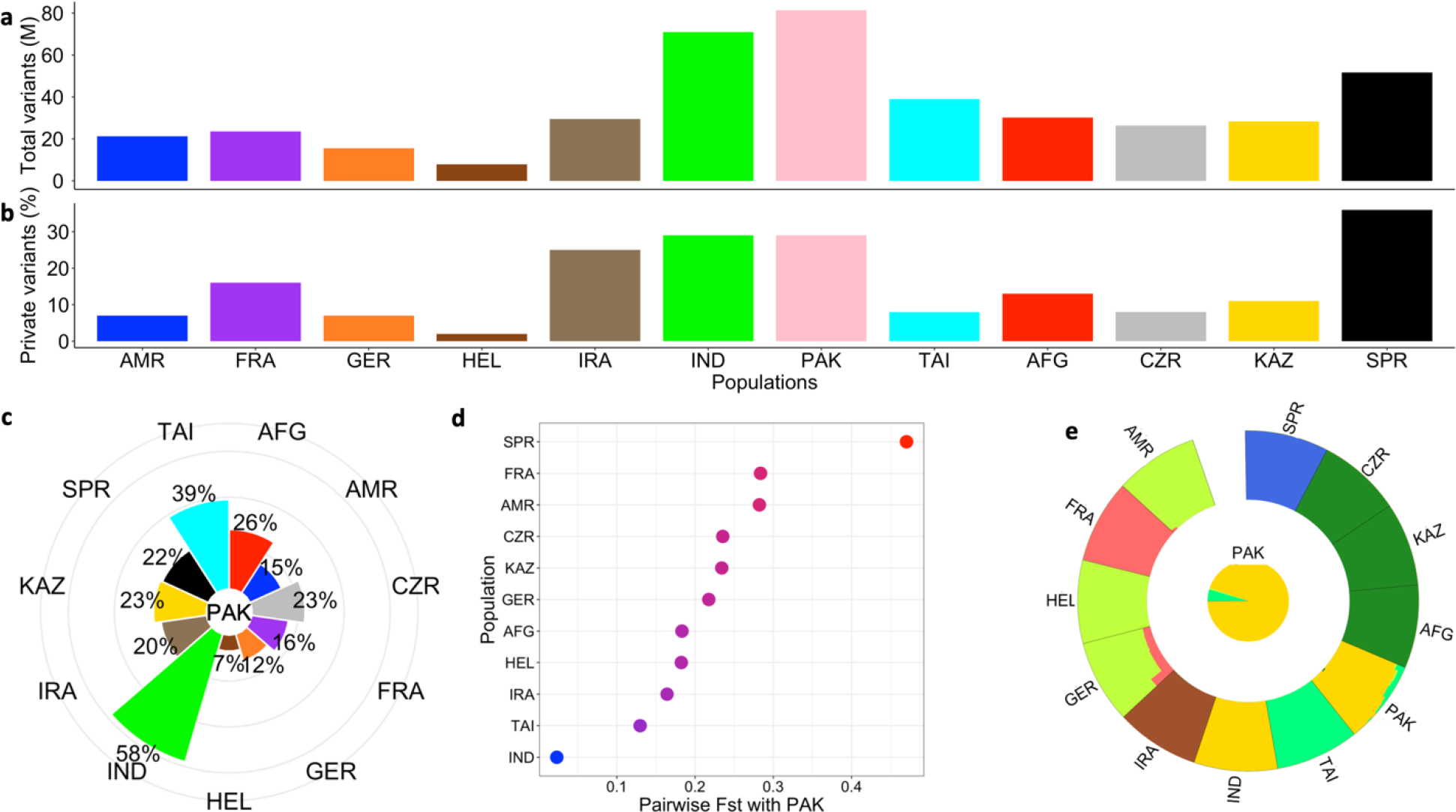
The landscape of genetic diversity across wild house mouse populations. (**a**) Total variants in each population (M = million) and (**b**) the proportion of private variants from (**a**). (**c**) Radial plot showing the proportion of segregating SNPs in each population that is shared with PAK. (**d**) Genome-wide estimates of pairwise F_st_ with PAK. (**e**) Global admixture proportion, with PAK highlighted in the middle to show high genetic affinity with the IND population.

Third, we expect that the genetic diversity in derived populations should largely subsample that present in ancestral populations. Consistent with this expectation, a phylogenetic tree constructed from autosomal SNPs places CAS (PAK and IND) at the base of the subspecies tree (Figure S1b).

Tests of these three predictions jointly support the conclusion that the ancestral homeland of *Mus musculus* is within the geographic boundary of modern-day India and Pakistan (Indo-Pakistan). Therefore, contemporary wild mouse populations from the Indo-Pakistan region could serve as proxies for the ancestral *M. musculus* population and will be broadly useful for the study of genetic diversity in modern house mice. We note that PAK and IND exhibit a higher proportion of shared genetic variants (Figure 2c) and greater genetic similarity (Figure 2d) than observed in other population comparisons. PAK and IND also exhibit a common admixture assignment (Figure 2e), confirming their high genetic affinity. This genetic similarity precludes confident designation of one population as a better representative of the ancestral population, although further wild mouse sampling from the Indo-Pakistan region could help resolve this question.

### Genetic differentiation between ancestral and non-ancestral house mice reveals legacy of genetic adaptation

As house mice radiated out of the ancestral geographic center in Indo-Pakistan, they encountered novel environmental conditions that likely imposed strong selective pressures. Contemporary wild mouse genomes likely bear signatures of historical adaptation, potentially manifesting as genomic regions with unusually high genetic differentiation between the ancestral Indo-Pakistani and non-ancestral house mouse populations. To uncover such signals, we computed F_ST_, a measure of relative genetic differentiation, in 50 kb windows across the genome for each ancestral-derived population pair. Overall, we observed strong, positive correlations between F_ST_ metrics computed for IND and each non-ancestral population and PAK and each non-ancestral population (Pearson’s correlation coefficient *r* ranges from 0.83 – 0.92, *p* < 0.001 Figure 3; Table S3). We focus on 73 unique genomic regions corresponding to the 1% of windows with the most extreme F_ST_. A GO analysis of genes within these regions reveals no statistically significant enrichment for genes annotated to specific biological processes, molecular functions, or cellular compartments. However, we note that these regions overlap multiple genes implicated in tumor growth (e.g. *Ttpal, Dhx36, Kdm5a, Nlrp1b, Uqcrb*), immune function (e.g. *Ankrd17, Trim30d, Nlrp1b*), and olfaction, biological processes enriched for signals of positive selection in prior wild mouse studies (Booker, et al. 2021; Lawal, et al. 2021).

**Figure 3:**
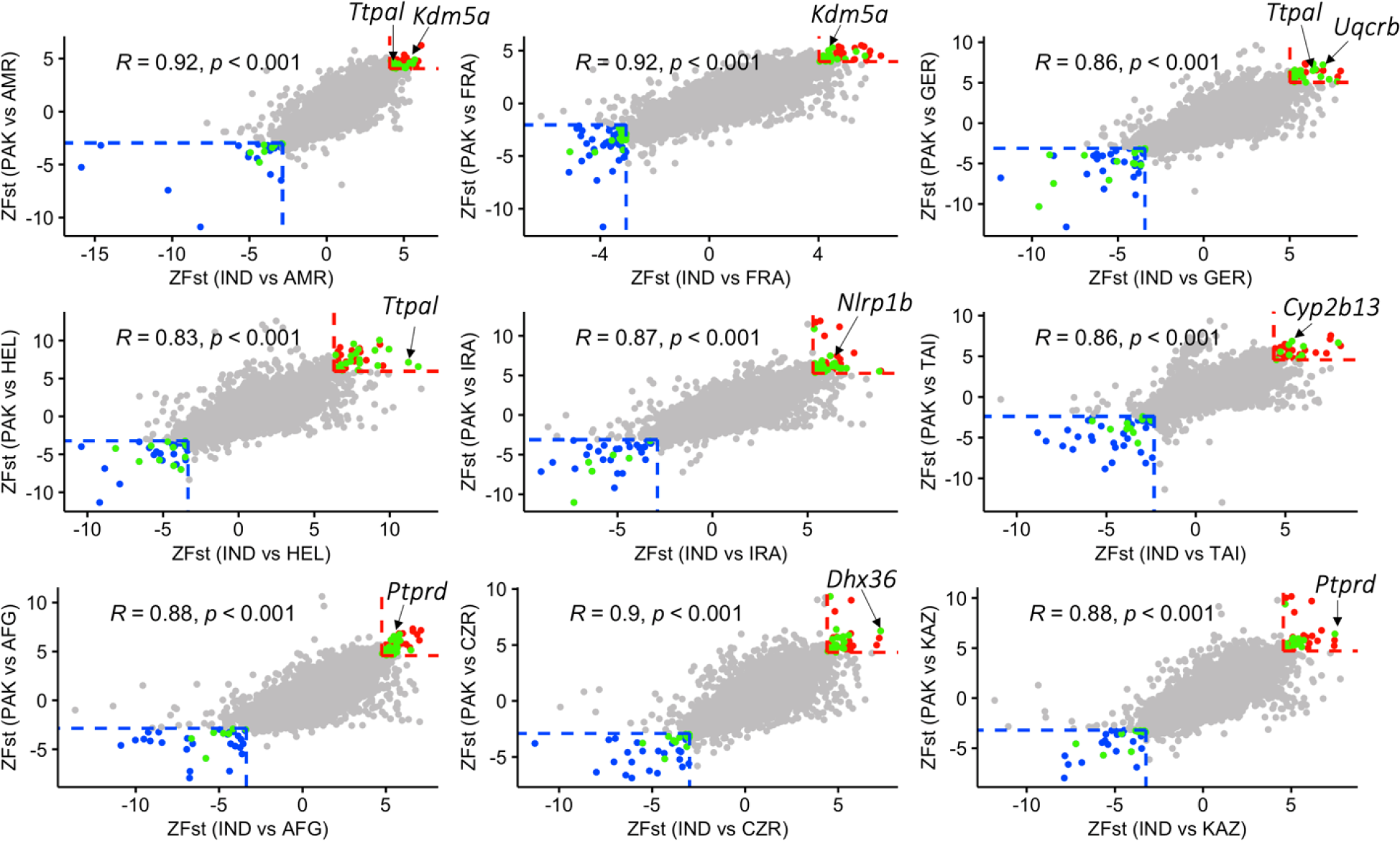
Dot plots depicting the relationship between genetic differentiation (normalized *F*_*st*_, *ZF*_*st*_) between each pair of ancestral (PAK/IND) and non-ancestral populations. The X-axis (Y-axis) shows *ZFst* between IND (PAK) and each of the non-ancestral populations. Red dots denote regions of high F_st_ in both ancestral-derived population comparisons and correspond to putative regions of adaptive evolution. The blue dots highlight regions of low F_st_ and represent putative targets of balancing selection. The green dots indicate regions containing known genes (see Table S3).

### Balancing selection has shaped the diversity of immune-related genes in wild house mice

In parallel to the analyses of population differentiation above, we performed a genome-wide scan for regions of unexpectedly high genetic similarity (low F_ST_) between ancestral and non-ancestral wild mouse populations. Such regions could represent (1) shared ancestral variation maintained via balancing selection or (2) recent introgression between ancestral and derived mouse populations. We used two approaches to differentiate between these possibilities.

Under balancing selection, alleles are maintained across lineages for longer time periods than expected, resulting in an excess of shared alleles between populations. To isolate regions maintained by balancing selection, we focus on regions with low F_ST_, but typical values of absolute divergence (D_XY_). While F_ST_ is reduced at sites evolving under balancing selection (Guerrero and Hahn 2017), D_XY_ tends to remain at background levels under balancing selection (Sicard, et al. 2015; Han, et al. 2017). In contrast, regions with low F_ST_ and high D_XY_ are likely sites where ancient haplotypes are incompletely sorted.

We plotted the relationship between F_ST_ × F_ST_ (Pearson’s correlation, 0.83 – 0.92, *p* < 0.001 Figure 3) and D_XY_× F_ST_ (*r* ranges from 0.3 – 0.38, *p* < 0.001 Figure S2) for each pair of ancestral and non-ancestral populations, flagging loci in the extreme lower 1% in each comparison. Using this approach, we identified a total of 57 candidate regions under balancing selection across the 11 *M. musculus* populations. An enrichment analysis on the 104 genes within these 57 regions identifies 48 GO terms and 25 KEGG pathways that are significantly over-represented. Remarkably, 31 (of 48) enriched GO terms and all 25 identified KEGG pathways are linked to biological processes associated with immune response (*p* < 0.05, Figure 4 and S3, Table S4). Further, at least 9 (of 31) immune-related GO terms are specifically linked to the major histocompatibility complex, a segment of the genome known to harbor immune-related genes subject to balancing selection (Croze, et al. 2016). These findings are consistent with the well-known role of balancing selection on genes with immune functions (Croze, et al. 2016; Koenig, et al. 2019), and suggest the likely phenotypic impact of ancestral variation on immune response and regulation in contemporary house mouse populations.

**Figure 4:**
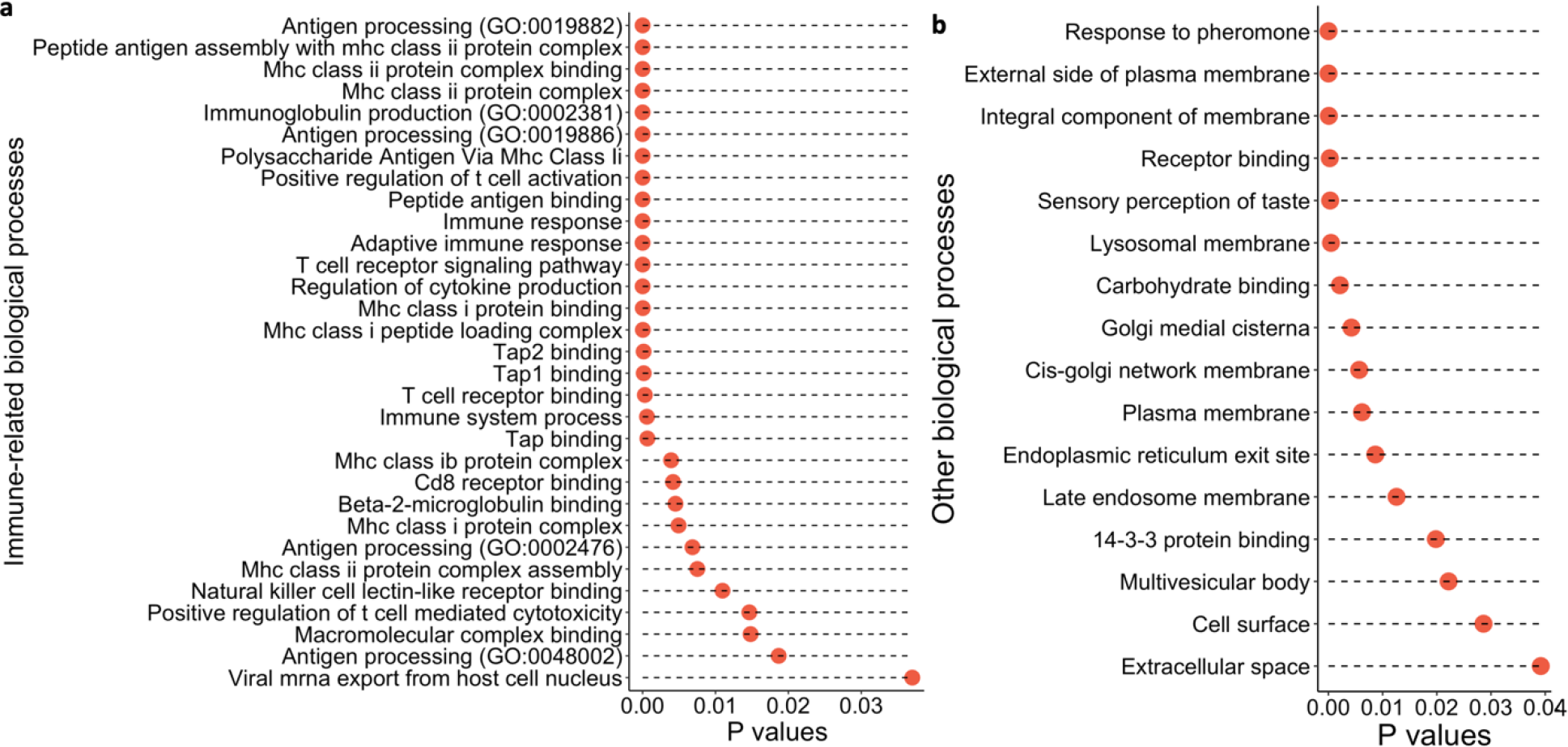
Regions under balancing selection are enriched for genes with immune-related biological functions **(a)** and non-immune related biological processes **(b)**. Immune related KEGG pathways identified from regions of balancing selection are presented in **Figure S3**.

**Figure S2:**
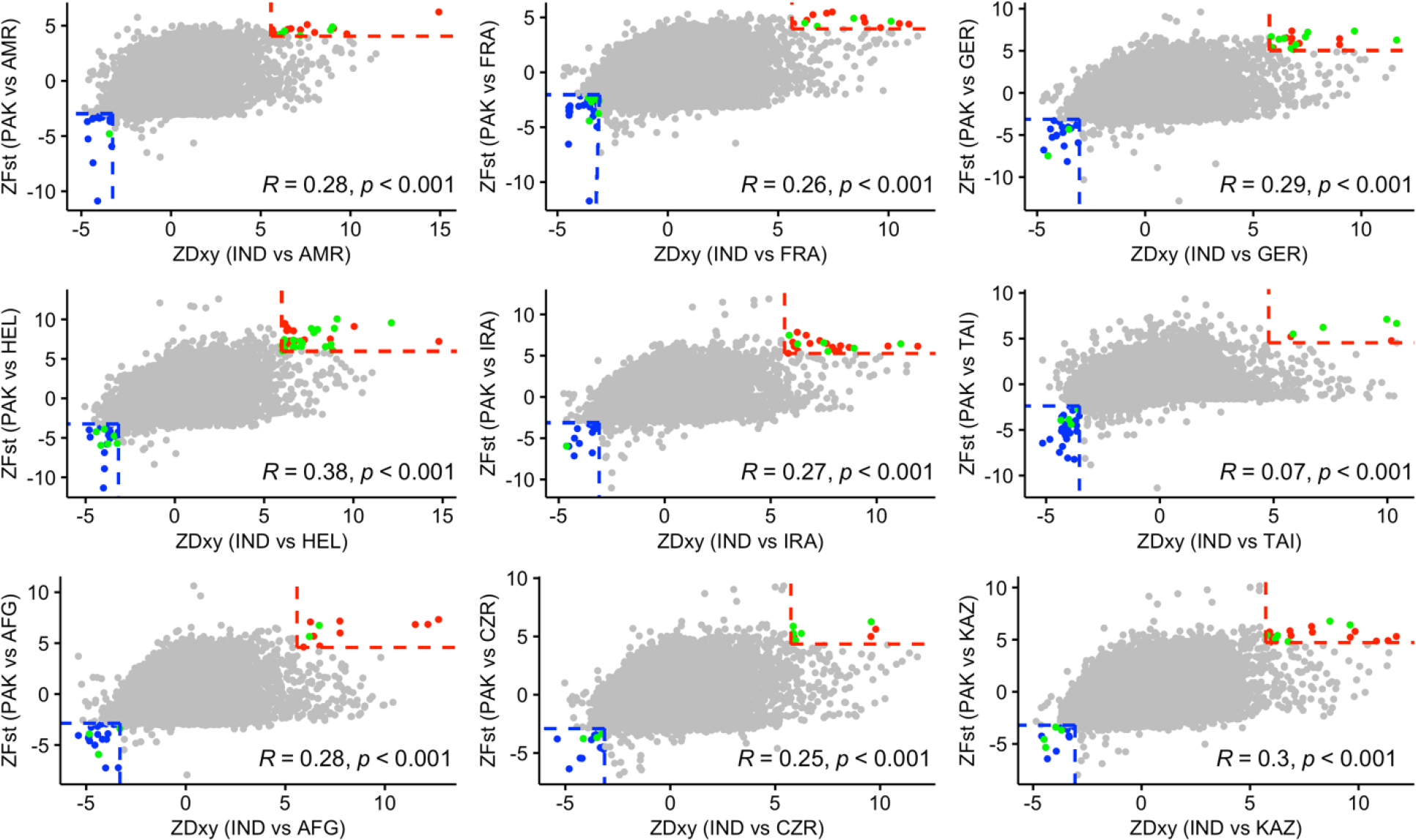
Dot plots of the relationship between F_st_ and D_xy_ in comparison between each pair of ancestral (PAK/IND) and non-ancestral populations. The X-axis (Y-axis) shows normalized D_xy_ between IND/PAK and each of the non-ancestral populations. Red dots denote regions of high F_st_ and D_xy_ in both ancestral-derived population comparisons and correspond to the 1% of regions with the most exceptionally high divergence. The blue dots are the 1% of regions with the lowest F_st_ and D_xy_ and correspond to regions potentially evolving under balancing selection. The green dots highlight regions containing known genes (see Table S4).

**Figure S3:**
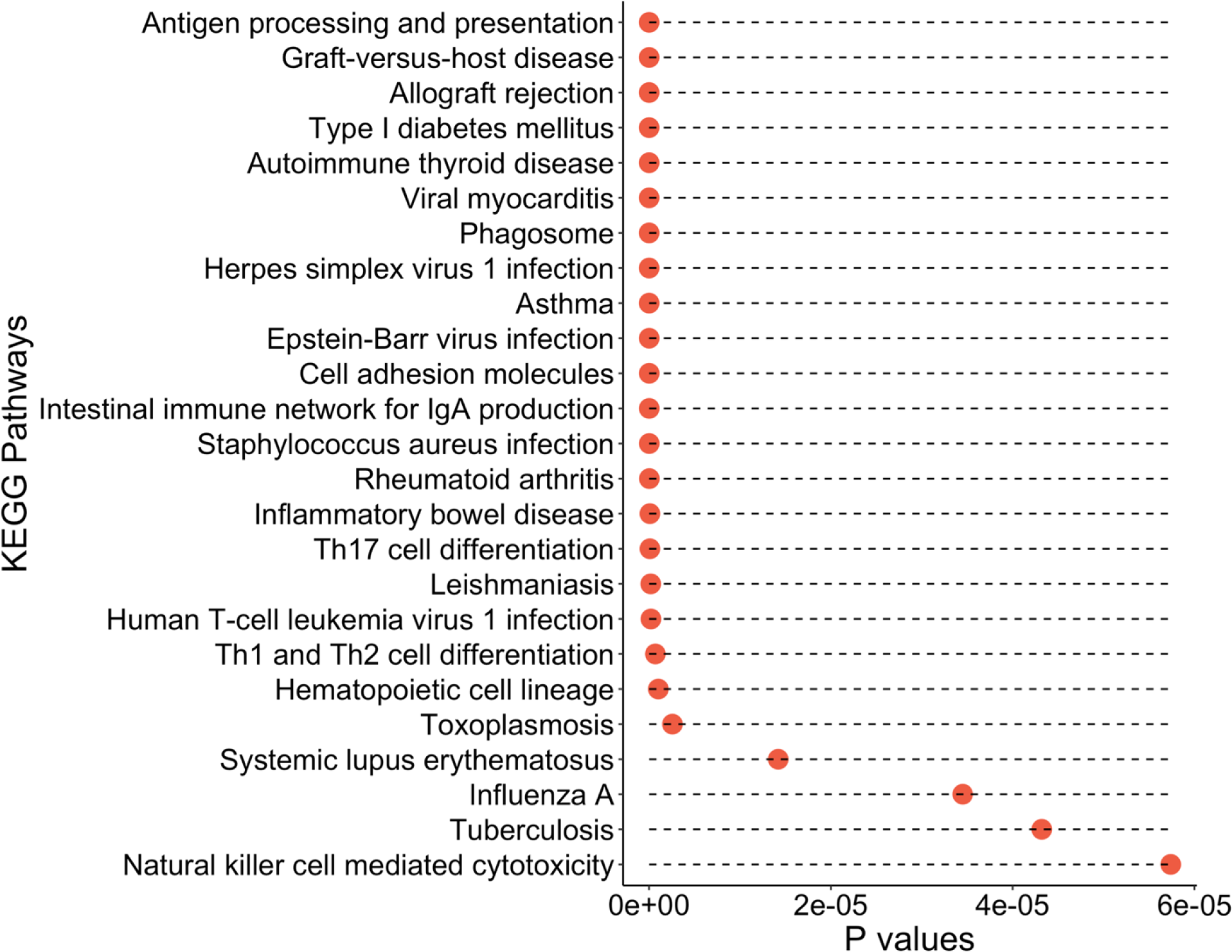
Regions under balancing selection harbor an enrichment of genes that function in immune-related KEGG pathways.

### Pervasive signals of incomplete lineage sorting across house mouse genomes

Genetic introgression between house mouse subspecies has recently been quantified on the genome-wide scale (*e*.*g*. (Ullrich, et al. 2017; Phifer-Rixey, et al. 2020; Fujiwara, et al. 2022)). Although house mouse subspecies are partially reproductively isolated (Turner and Harr 2014), these studies have concluded that subspecies hybridization has left substantial footprints in the genomes of wild mice. This result is somewhat surprising, given that partial reproductive isolation between subspecies tends to reduce introgression rates (Zhou, et al. 2017). Incomplete lineage sorting (ILS) is defined by the retention of ancestral polymorphisms in two or more descendent taxa and can mimic the genomic signal of recent introgression events (Maddison 1997). The relatively young evolutionary history of house mouse subspecies raises the possibility that previously reported signals of subspecies hybridization may simply represent regions of the genome with deep coalescence times that pre-date the divergence between subspecies. Using our comprehensive dataset of ancestral mouse genomes, we explore this alternative hypothesis using multiple, complementary approaches.

First, we estimated genome-wide levels of admixture between the ancestral Indo-Pakistan population and derived populations of DOM, MUS, and CAS using Patterson’s *D* and *f*4 statistics (Green, et al. 2010; Durand, et al. 2011; Patterson, et al. 2012). We observe the strongest admixture signals between the Indo-Pakistan CAS populations and the Taiwanese population of CAS, presumably reflecting the recent origin of the TAI population. The degree of admixture was significantly higher between CAS and MUS than between CAS and DOM, although there is variation in the degree of CAS admixture across DOM populations (Figure S4). Overall, findings from these analyses align with previously reported subspecies relationships (White, et al. 2009; Phifer-Rixey, et al. 2020; Lawal, et al. 2022), providing additional support for CAS and MUS as sister taxa.

**Figure S4:**
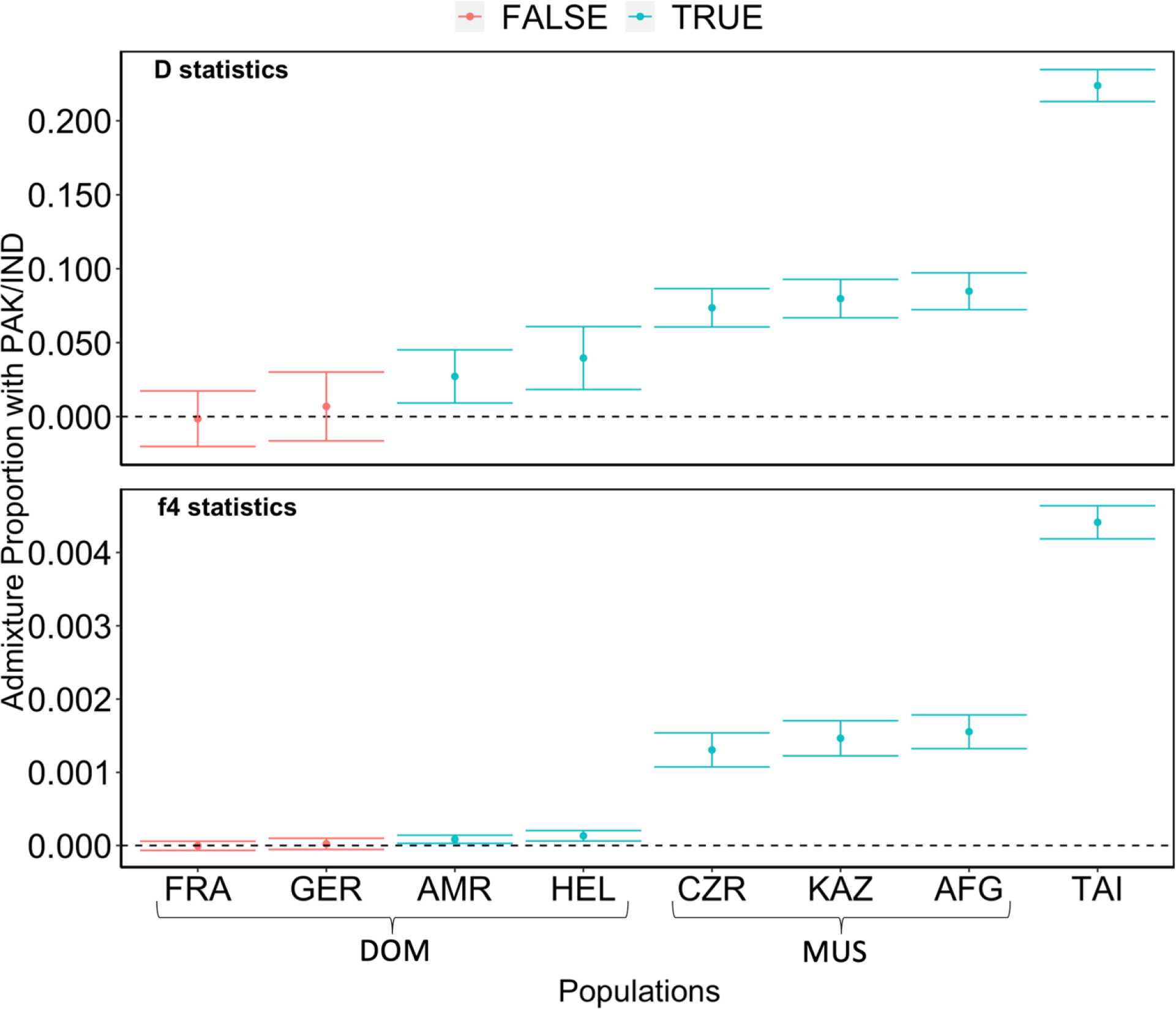
Genetic admixture between the ancestral Indo-Pakistan CAS populations and other mouse populations. *D* or *f*4 statistics that do not significantly deviate from zero comply with the null hypothesis of no admixture with the ancestral population and are color-coded as “FALSE”. Populations with significantly non-zero D or f4 statistics indicating an excess of shared derived alleles with the ancestral Indo-Pakistan populations are color-coded “TRUE”. Such admixture signals could be due to introgression with the Indo-Pakistan populations or incomplete lineage sorting. The IRA population was used to polarize the admixture signal with the outgroup and was therefore omitted from this analysis.

The significant departure of *D* and *f*4 from zero reveals an excess of shared derived alleles between the Indo-Pakistan CAS populations and most other wild mouse populations at the whole-genome scale (Figure S4). These signals could arise from ILS or represent pervasive, recent introgression (Green, et al. 2010; Durand, et al. 2011). To distinguish between these two possibilities and identify specific loci impacted by introgression, we quantified locus-specific admixture proportions using *f*_d_, a bidirectional introgression statistic designed on the basis of the four-taxon ABBA-BABA test (Martin, et al. 2015). We then complement this approach with manual analysis of locus-specific phylogenetic trees. Our expectation is that introgression will alter the topology of the previously established subspecies tree (White, et al. 2009; Phifer-Rixey, et al. 2020; Lawal, et al. 2022), whereas ILS will distribute alleles more or less randomly across samples, irrespective of their associated subspecies or population identity.

For each wild mouse population, we identified the 10 regions with the most extreme *f*_d_ statistic using alleles in both IND and PAK as proxy ancestors (see Methods). After merging adjacent windows and identifying regions common across populations, this effort flagged a total of nine unique genomic windows (Figure S5). Eight of these nine regions exhibit high *f*_d_ signals in multiple populations, suggesting widespread allele sharing across taxa (Figure S5). Phylogenetic analyses failed to offer conclusive evidence of introgression at any of these eight regions. Overall, samples are not strictly clustered according to the established subspecies tree (White, et al. 2009; Keane, et al. 2011; Phifer-Rixey, et al. 2020; Fujiwara, et al. 2022; Lawal, et al. 2022), with significant interspersion of samples from different subspecies assignments (Figure S6). We interpret these observations as most consistent with ILS, rather than introgression.

The remaining *f*_d_ outlier region is specific to analyses with the KAZ population (chr2: 86.62-86.71 Mb). This region overlaps a cluster of olfactory receptors, which are highly copy number polymorphic and sequence-variable across house mice (Pezer, et al. 2015). While the associated phylogenetic tree for this locus suggests a possible history of introgression from IND and/or PAK into KAZ (Figure S6i), we cannot rule out the alternative interpretation that reference biases and mapping errors drive an artificial signal of introgression at this locus.

Overall, we find little evidence for historical introgression from the Indo-Pakistan ancestral population into modern-day mouse populations, suggesting limited gene flow out of the ancestral homeland in recent time. Instead, we favor the interpretation that genome-wide departures from expected levels of allele sharing between our surveyed populations reveal pervasive footprints of ILS in contemporary house mouse genomes.

**Figure S5:**
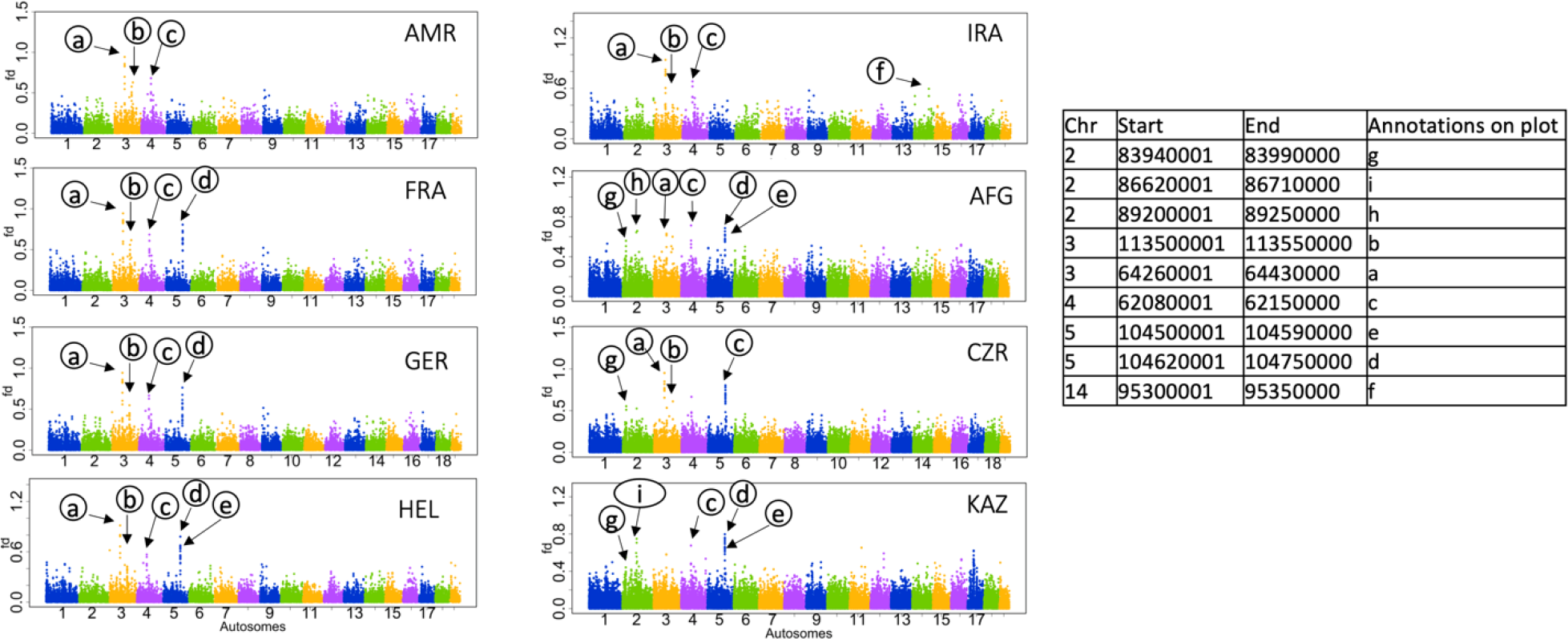
Manhattan plot showing the distribution of locus-specific admixture proportion (*f*_d_) between ancestral and derived house mouse populations (see Methods). *f*_d_ ranges from 0 (no admixture) to 1 (absolute admixture), with each dot representing the admixture proportion estimated across a 50kb window. The top ten windows were analyzed in each population. Following the merger of adjacent windows and integration across populations, this corresponded to a total of nine unique regions, with coordinates indicated in the table. These outlier regions are annotated (a) to (i) on the plots, with most regions shared by multiple populations. Region (i) is the only *f*_d_ outlier unique to a single population (KAZ).

**Figure S6:**
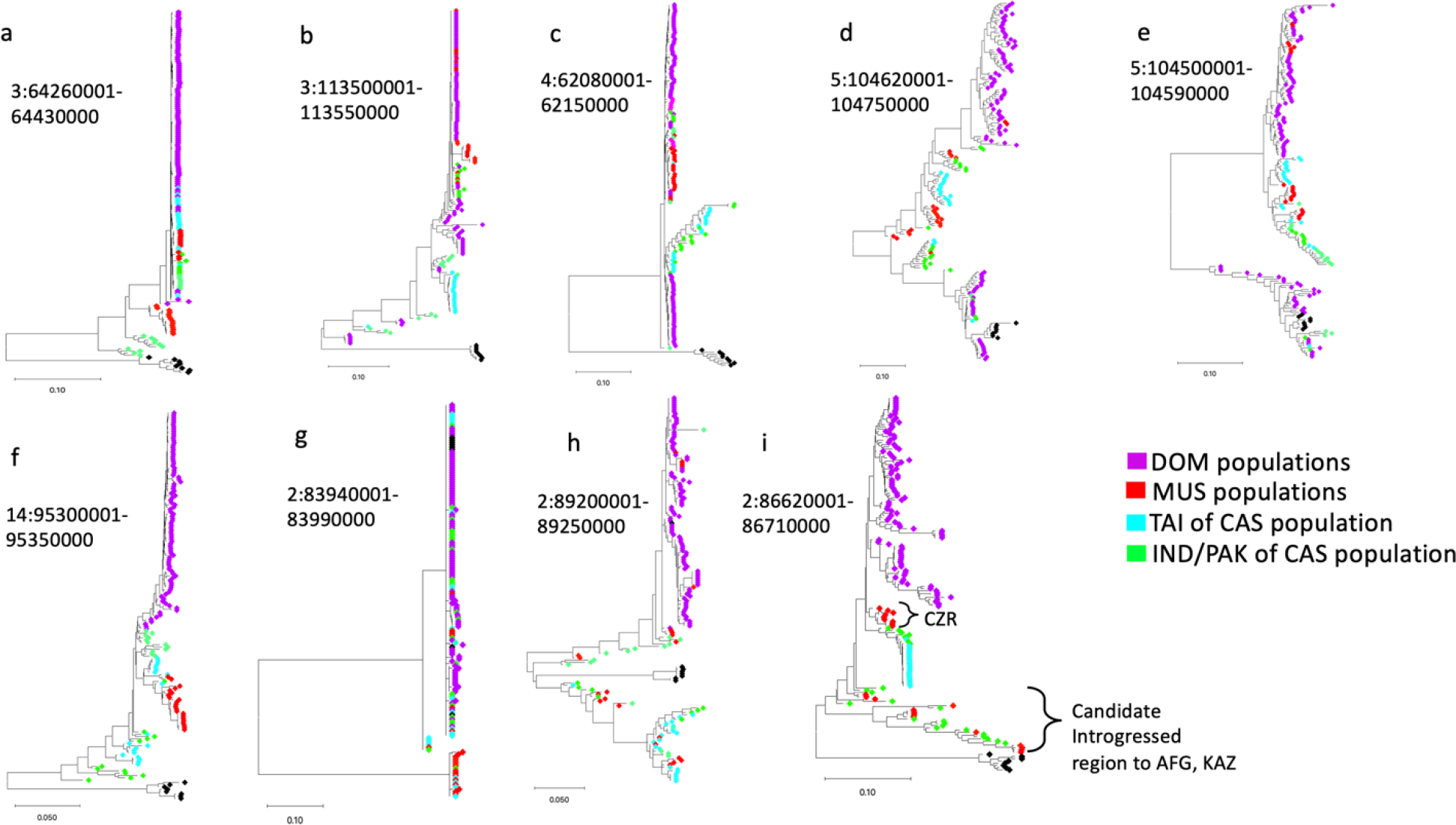
Phylogenetic trees for the regions identified in Figure S5. Samples are color-coded by their respective subspecies. TAI, a derived population of CAS, is assigned a unique color to distinguish it from the ancestral IND/PAK populations of CAS.

## DISCUSSION

Despite the extensive genomic resources and deep understanding of genetic diversity in inbred laboratory mouse strains, little is known about the organization of genetic variation in wild house mouse populations or the population genetic forces that contribute to this variation. Here, we analyzed 169 wild mouse genomes – to our knowledge, the largest dataset compiled thus far – to define key demographic events during house mouse evolution, resolve the geographic ancestry of house mouse subspecies, and highlight the contribution of ancestral variants to contemporary patterns of diversity in house mouse populations across the globe. Our work leverages the power of large population genomic datasets to provide important new insights into the natural evolutionary history of this important model organism.

We provide strong support for the hypothesis that house mice emerged in the region of modern-day India and Pakistan. Mice from this region exhibit greater effective population sizes and harbor increased genetic diversity relative to mice from other populations. Further, genetic variation in this region largely subsumes that present in other geographic regions, as expected if subsequent migratory waves only subsampled alleles segregating in these ancestral populations. Overall, these trends bear clear parallels to the patterns of genomic diversity within and outside of the ancestral homelands of many other species, including humans (Campbell and Tishkoff 2008; Chan, et al. 2019).

In addition, our work provides key estimates for major events on the house mouse evolutionary timeline. Prior subspecies divergence estimates suggested near-simultaneous radiation of the three cardinal house mouse subspecies (*M. m. castaneus, M. m. domesticus*, and *M. m. musculus*) ∼150 - 500 kya (Boursot, et al. 1996; Geraldes, et al. 2008; Geraldes, et al. 2011; Bonhomme and Searle 2012; Suzuki, et al. 2013; Phifer-Rixey, et al. 2020). Our findings indicate an initial emergence of an ancestral *M. m. castaneus* population ∼700 kya, slightly deeper in time than other published estimates. *M. m. domesticus* subsequently diverged ∼360 kya, followed by *M. m. musculus* ∼256 kya. These latter two estimates are slightly older than those presented in some recent reports (Phifer-Rixey, et al. 2020; Fujiwara, et al. 2022). However, we acknowledge that reported divergence times are highly sensitive to assumptions about generation time, a parameter that remains poorly understood for wild mice and that likely varies between house mouse subspecies and populations in nature. Here, we have assumed 1 generation/year in all populations, a value equal to that employed in other studies yielding distinct estimates (e.g., Fujiwara, et al. (2022)). Methodological differences undoubtedly also contribute to differences in subspecies divergence estimates across studies. In particular, prior analyses have invoked approaches known to produce unreliable estimates of young divergence times (Terhorst, et al. 2017; Fujiwara, et al. 2022), relied on genotypes at a limited number of loci that may be subjected to ascertainment bias (Geraldes, et al. 2008; Geraldes, et al. 2011; Suzuki, et al. 2013), or the use of model-based approaches that may be sensitive to misspecification of demographic history (Geraldes, et al. 2008; Phifer-Rixey, et al. 2020). Importantly, our estimates are free of these limitations, leveraging whole genome data from multiple samples and the assumption-free coalescent framework implemented in SMC++.

Our findings reinforce the current model of house mouse demographic history and global colonization suggested by prior studies (Bonhomme and Searle 2012; Didion and de Villena 2013). Mice from an ancestral population within the Indo-Pakistan migrated southeastward into modern-day Iran around 360 kya, giving rise to *M. m. domesticus*. Subsequent migratory events facilitated by human movements led to the expansion of *M. m. domesticus* out of the Fertile Crescent into Western Europe, the Mediterranean basin, and Northern Africa. Over the last few hundred years, human seafaring traffic enabled *M. m. domesticus* to colonize North and South America, New Zealand, and Australia (Gabriel, et al. 2011; Agwamba and Nachman 2023). A potential second migratory wave out of Indo-Pakistan around ∼260 kya followed a more northern trajectory into modern-day Afghanistan, yielding *M. m. musculus*. Mice from this migratory episode subsequently experienced range expansion into Eastern and Central Europe and Northern Asia (Prager, et al. 1998). However, some genetic and archeological evidence supports the emergence of *M. m. musculus* from an initial source population in Transcaucasia (Duvaux, et al. 2011). Due to limited sampling, the origins of this subspecies remain contentious, and additional phylogeographic analysis of wild-caught *M. m. musculus* mice from across Eurasia will be required to fully clarify the origin of this subspecies. A third migration wave, likely occurring around 460 kya led to the expansion of house mice out of the ancestral geographic center in the Indo-Pakistan valley into Southern Asia (Suzuki, et al. 2013). Future work is needed to refine the dates of later migration episodes outside the Indo-Iranian valley, but it is widely believed that these dispersions were tied to the movements of early human agricultural societies in the Neolithic age. Indeed, the geographic regions that encompass modern-day Iran and Afghanistan were home to some of the earliest farmers (Broushaki, et al. 2016).

Following migration out of the ancestral Indo-Pakistan homeland, house mice were exposed to new environments that undoubtedly imposed new and intense selection pressures. The ability of house mice to rapidly adapt to these new conditions was paramount to their successful global colonization. Although prior studies have identified genome-wide targets of positive selection in house mice (Staubach, et al. 2012; Phifer-Rixey, et al. 2018; Booker, et al. 2021; Lawal, et al. 2021), focus has been mostly restricted to the discovery of selection on new mutations, rather than selection on standing variation. The waiting time to new adaptive mutations may be substantial, implying that adaptation may often be driven by selection on standing variants (Duvaux, et al. 2011; Reid, et al. 2016; Lai, et al. 2019). We explored the possibility that ancestral alleles provide an important source of adaptive variation in derived house mouse populations by identifying genomic regions with unusually large allele frequency shifts in comparisons with ancestral IND/PAK (Figure 3). This molecular signature is expected if selection acts to increase (or decrease) the frequency of a given allele (and linked sites) due to their adaptive benefit in a new environment. While our use of a heuristic cutoff does not allow for precise estimation of the proportion of adaptation mediated by standing versus novel variation, our findings nonetheless emphasize a key contribution of ancestral variation to the legacy of adaptive evolution in wild house mice.

We also spotlight several regions of unexpectedly low genetic differentiation between ancestral and non-ancestral populations. Such regions harbor alleles that coalesced prior to subspecies divergence and represent candidate targets of historical balancing selection. Intriguingly, virtually all of the genes within these candidate regions have immune-related functions. This finding adds to a deep body of empirical literature demonstrating the importance of balancing selection for the long-term maintenance of alleles in immune-related genes (Andrés, et al. 2009; Koenig, et al. 2019; Minias and Vinkler 2022).

Our work also helps define the extent to which ancestral house mouse variants contribute to genetic diversity in derived populations through both gene flow and incomplete lineage sorting. House mice hybridize at regular zones of secondary contact in the wild, leading to both stable hybrid populations (Yonekawa, et al. 1988) and more restricted signatures of admixture due to partial reproductive isolation (Turner and Harr 2014). Prior studies have also suggested extensive population structure on local geographic scales (Morgan, et al. 2022), but little evidence for persistent gene flow between populations (Ullrich, et al. 2017; Phifer-Rixey, et al. 2020; Fujiwara, et al. 2022). To our knowledge, our investigations present the first genome-wide tests of gene flow from ancestral mouse populations into derived populations. Overall, our findings indicate that regions of shared ancestry in the analyzed wild mouse genomes are more likely due to ILS than introgression. However, our surveyed populations are geographically isolated, and future analyses featuring additional genomic sequences from populations within or near the ancestral region may reveal signatures of contemporary or recent introgression. Regardless, we conclude that the recent emergence of *M. musculus* has offered insufficient time for many alleles to fix along independent subspecies lineages, leading to significant phylogenetic discordance across the house mouse genome, a finding in line with observations from prior work (White, et al. 2009).

## Conclusions

Our population genomic analyses of wild house mice from multiple populations and subspecies enumerate key events in the evolutionary history of this important model species. We show that *M. musculus* emerged ∼700 kya in the area of modern India and Pakistan, with two subsequent radiations ∼360 and ∼260 kya giving rise to the *M. m. domesticus* and *M. m. musculus* subspecies, respectively. Our findings reinforce the hypothesis that *M. m. musculus* and *M. m. castaneus* are sister taxa, bringing a larger dataset and greater representation of genomes from the ancestral homeland region to bear on this question than earlier studies. We also show that ancestral alleles have provided the raw material for house mouse adaptive evolution outside the ancestral home range and, in some cases, have been maintained by adaptive processes for exceptionally long times. Finally, we demonstrate significant sharing of ancestral alleles across house mouse subspecies due to incomplete lineage sorting, with little evidence for introgression among our studied populations. Overall, our findings refine current understanding about the geographic origins, demographic history, and population genetic processes that impact global house mouse diversity. This knowledge bears on the ultimate sources of the genetic variation that fuels biomedical discovery in laboratory mice.

## Materials and Methods Published genomes

We retrieved 169 previously published genomes belonging to *M. m. domesticus* (America, AMR = 50, France, FRA = 28, Germany, GER = 7, Heligoland, HEL = 3, Iran, IRA = 7), *M. m. castaneus* (India, IND = 10, Pakistan, PAK = 14, Taiwan, TAI = 20), *M. m. musculus* (Afghanistan, AFG = 6; Czech Republic, CZR = 8; Kazakhstan, KAZ = 8), and *M. spretus* (Spain, SPR = 8) (Davies 2015; Harr, et al. 2016; Phifer-Rixey, et al. 2018; Lawal, et al. 2022). PAK genomes (n=14) are available on the NCBI Short Read Archive under the BioProject accession PRJNA851025 (https://www.ncbi.nlm.nih.gov/sra/PRJNA851025).

### Sequence alignment and variant calling

Reads were mapped to the mm10 reference genome using the default parameters in BWA version 0.7.15 (Li 2013). We followed the standard Genome Analysis Toolkit (GATK; version 4.2) pipeline for subsequent pre-processing prior to variant calling (Li and Durbin 2010; Auwera, et al. 2013). Variant calling was performed individually for each sample using the “-ERC GVCF” mode in the “HaplotypeCaller”. Samples were then jointly genotyped using the “GenotypeGVCFs” GATK function and trained with previously ascertained mouse variants (Keane, et al. 2011) using both the “VariantRecalibrator” and “ApplyVQSR” option of GATK. For the latter, the truth sensitivity level to initiate filtration was set to the default (i.e., 99). Our downstream analysis includes only biallelic single nucleotide variants.

### Demographic history inference

We inferred the demographic histories of each surveyed wild mouse using SMC++ v1.15.2 (Terhorst, et al. 2017). SMC++ uses the sequentially Markovian coalescent to infer past population sizes and estimate timing of population size shifts. Although multiple methods for demographic inference are available, we used SMC++ for several reasons. First, SMC++ utilizes several genomes at the same time, in contrast to the alternative programs (*e*.*g*. PMSC (Hey, et al. 2018)) that are restricted to single or limited numbers of individuals (Beichman, et al. 2017). Second, SMC++ is computationally efficient and does not require phased data (Terhorst, et al. 2017). Third, SMC++ improves accuracy for the estimation of recent population sizes relative to other programs (Mather, et al. 2020).

A total of 89 (of 161) house mouse genomes were used for estimating subspecies demographic history. Due to recent colonization events in TAI and HEL, and limited genome coverage in AMR, these populations were excluded from subspecies-level analysis. However, for population level inferences, we included genomes from all available populations and samples (n=161).

SNPs from each chromosome were converted into the SMC++ input format using the “vcf2smc” program supplied with the SMC++ release. Effective population sizes were then estimated over time for each population using the “estimate” option. We specified a per-generation mutation rate of 5.4×10^-9^ (Uchimura, et al. 2015), piecewise spline, and invoked defaults for other options. To convert divergence times to years, we assumed one generation per year. The “plot” subcommand of SMC++ was used to generate an output CSV file which was then viewed using the ggplot2 package in R (Wickham 2016).

### Identifying ancestral house mouse populations

We collected multiple population genetic summary statistics to facilitate inference of the ancestral population(s) of house mouse. First, we estimated nucleotide diversity in 100kb windows (50kb slide) using VCFtools v0.1.16. We used estimates of effective population size from SMC++, described above. To identify population specific variants, we first used VCFtools v0.1.16 to partition the comprehensive VCF file into per-population files. VCFtools was then used to filter variants on each population-level VCF file using the following options: “--maf 0.001 --max-maf 0.999 --max-missing 0.9”. Per-population VCF files were then intersected using BCFtools isec, invoking the “--complement” option (Li, et al. 2009).

Admixture was assessed using ADMIXTURE v1.3.0 software (Alexander, et al. 2009), with ten independently replicated runs indicating convergence on consistent cluster assignments. EvalAdmix was used to assess the goodness of fit of ancestry assignments (Garcia-Erill and Albrechtsen 2019), with the best fitting cluster assignment plotted using ancestryPainter (Feng, et al. 2018).

To construct maximum likelihood tree, we first thinned the comprehensive VCF file to 656,538 SNPs using VCFtools v0.1.16 with flags “--max-missing 1 --thin 1000”. Molecular evolutionary model inference was performed using jModeltest 2.1.7 (Darriba, et al. 2012), with Phyml 3.0 used to compute the approximate likelihood ratio score for each branch based on the best predicted model (Guindon and Gascuel 2003). The resulting tree was viewed in MEGA version 11.0.11 (Stecher, et al. 2020).

### Genetic differentiation analysis

We compute the fixation index, F_ST_, between all pairs of non-ancestral and ancestral populations in 50kb windows (20kb slide) using the script “popgenWindows.py” (available at https://github.com/simonhmartin/genomics_general (Martin, et al. 2015)). F_ST_ values were normalized to Z-scores and then compared and plotted between each population pair using “ggscatter” in ggplot (Wickham 2016). We focus on the extreme 1% of points with the highest and lowest Pearson correlation coefficient.

### Identifying genomic signals of balancing selection

Balancing selection can result in the maintenance of shared ancestral polymorphism between divergent populations. Under such a scenario, F_ST_ is reduced (Guerrero and Hahn 2017), but absolute divergence D_XY_ is often unchanged or elevated (Sicard, et al. 2015; Han, et al. 2017). To identify putative targets of balancing selection in wild house mouse genomes, we identified the 1% lower tail of the distribution of F_st_ X F_st_ correlations between PAK/IND and non-ancestral populations. We then intersected these regions with D_XY_ measures and restrict our attention to those regions with D_XY_ values equal to or elevated relative to genome wide D_XY_.

### Identification of introgressed regions between house mouse subspecies

We quantified genome-wide admixture proportions using multiple methods. First, we used the “admixr” package (Petr, et al. 2019) to compute *D* and *f*4 statistics (Durand, et al. 2011). These statistics allow for the detection of excess admixture or an enrichment of shared derived alleles in DOM and MUS populations relative to PAK/IND. Admixture was quantified using *M. spretus* SPR as an outgroup. To detect specific loci with an excess of admixture, we used *f*_*d*_, a bidirectional introgression approach designed on the basis of the four taxon ABBA-BABA test (Martin, et al. 2015). We defined our three in-groups as *P*_1_ (either PAK or IND), *P*_2_ (each of the non-ancestral house mouse populations), *P*_3_ (either PAK or IND). *Mus spretus* was used as the outgroup. If *P*_1_ was set to IND, then *P*_3_ was set to PAK, and *vice versa*. We differentiated a signature of ILS from that of introgression by comparing *f*_*d*_ and D_XY_. Both statistics were computed over 50kb windows (20 kb slide), with windows containing fewer than 100 SNPs excluded. We selected the top 10 windows in each population and collapsed adjacent windows into single contiguous regions using BCFtools merge. A phylogenetic tree was then constructed for each region using jModelTest and phyml as described above. We expect that regions subject to introgression will cluster a subset of individuals from the source and recipient populations. We invoked this expectation to confirm regions of introgression and infer the directionality of introgression events.

### Functional annotation

Genes within windows identified as putative targets of positive selection and balancing selection were retrieved using Ensembl BioMart version 102 (Kinsella, et al. 2011). These gene lists were then input into the Database for Annotation, Visualization, and Integrated Discovery (DAVID version 6.8) to test for GO term and KEGG pathway enrichment (Huang, et al. 2009). All genes in the *M. musculus* genome were used as background. Overrepresented gene clusters were identified by Fisher’s Exact tests (*p* < 0.05) and visualized with ggplot2 (Wickham 2016).

## Funding

RAL was supported by a JAX Scholar Award from The Jackson Laboratory. BLD is supported by a Maximizing Investigators Research Award from the National Institutes of Health (R35 GM133415).

## Declaration

Both authors declare no competing interests.

## Notes

### Competing Interest Statement

The authors have declared no competing interest.

